# De novo design of RNA and nucleoprotein complexes

**DOI:** 10.1101/2025.10.01.679929

**Authors:** Andrew Favor, Riley Quijano, Elizaveta Chernova, Andrew Kubaney, Connor Weidle, Morgan A. Esler, Lilian McHugh, Ann Carr, Yang Hsia, David Juergens, Kenneth D. Carr, Paul T. Kim, Yuliya Politanska, Enisha Sehgal, Paul S. Kwon, Robert J. Pecoraro, Cameron Glasscock, Andrew J. Borst, Frank DiMaio, Barry L. Stoddard, David Baker

## Abstract

Nucleic acids fold into sequence-dependent tertiary structures and carry out diverse biological functions, much like proteins. However, while considerable advances have been made in the *de novo* design of protein structure and function, the same has not yet been achieved for RNA tertiary structures of similar intricacy. Here, we describe a generative diffusion framework, *RFDpoly*, for generalized *de novo* biopolymer (RNA, DNA and protein) design, and use it to create diverse and designable RNA structures. We design RNA structures with novel folds and experimentally validate them using a combination of chemical footprinting (SHAPE-seq) and electron microscopy. We further use this approach to design protein-nucleic acid assemblies; the crystal structure of one such design is nearly identical to the design model. This work demonstrates that the principles of structure-based *de novo* protein design can be extended to nucleic acids, opening the door to creating a wide range of new RNA structures and protein-nucleic acid complexes.

## Introduction

Noncoding RNAs adopt complex three-dimensional structures that underpin a wide range of biological functions, including genetic regulation, catalysis, and scaffolding of biochemical machinery. In the field of nucleic acid design, most strategies have focused on three main areas: aptamer generation via randomized sequence selection^1,2^, the design of small RNAs with defined secondary structures^3,4^, and the construction of large, geometrically regular RNA/DNA origami^5,6,7,8^. Aptamer generation has been largely structure-independent, while secondary structure-based approaches focus on two-dimensional base pair templates and do not account for three-dimensional geometry, which limits their utility in applications requiring precise spatial control. Three-dimensional structure design methods—such as nucleic acid origami, hierarchical motif alignments^9^, and parametric design based on propagation of idealized geometry^10,11^— typically rely on rigid geometric assumptions, enforcing uniform helical axes and simplifying the placement of structural features like crossovers and base contacts. While these approaches can generate sophisticated and functional nucleic acid structures, they constrain the accessible design space and fail to capture the complexity observed in native RNA folds, which often feature multi-angled junctions, bulges, bent helices, and loop-loop contacts, which are likely important for more complex functions, such as catalysis. Thus, there remains a critical need for structure-oriented nucleic acid design approaches that enable exploration of the space of diverse three-dimensional structures exemplified by natural RNAs.

We reasoned that recent advances in generative deep learning for protein design could be leveraged to address the current limitations in nucleic acid design. Generative models such as RFdiffusion have shown considerable success in structure-oriented protein design^12,13,14^, and the RoseTTAfold framework has been adapted for nucleic acid structure prediction^15,16^. Although denoising diffusion probabilistic models (DDPMs) have been applied to generate short RNA monomers^17,18,19,20,21^, to date, these efforts lack hierarchical design capabilities, lack multi-polymer structure generation, and none have yet been experimentally validated. We set out to generalize the RFdiffusion *de novo* protein design approach to nucleic acids, and to explore the use of this method by designing novel RNA structures and protein-nucleic acid assemblies.

### Computational approach

Our design approach proceeds in two steps. First, we generate RNA or DNA backbone structures using an extended version of RFdiffusion (*RFDpoly*), described in the following paragraphs. Second, we design base sequences on the generated backbones using NA-MPNN^22^, which generalizes the widely used ProteinMPNN^23,24^ to nucleic acids.

RFDpoly extends RFdiffusion to enable the generation of nucleic acid structures by building on RF2-AllAtom^15,16^, which models both proteins and nucleic acids and accepts conditional information through 1D, 2D, and 3D channels. Previous RFdiffusion implementations could only denoise protein structures around fixed nucleic acid ligands; RFDpoly generates nucleic acid structures as well. Unlike proteins, where all non-frame backbone atoms (O, C_β_) can be deterministically placed given frame coordinates, nucleic acid backbones have increased conformational freedom due to variable sugar and phosphate geometries. To address this, RFDpoly still denoises frame rotations and translations as in previous RFdiffusion models, but goes further by using rigid body parameters *and* torsion angles predicted from RoseTTAfold to construct complete nucleic acid backbones (and side chains, when sequence or motif geometry is provided). To increase the extent to which the denoised frames determine nucleobase positioning, we switched from the phosphate-centered frame atoms used in RF2-AllAtom (OP1-P-OP2) to O4’-C1’-C2’ frame atoms, which are closer to bases.

To condition the diffusion process, RFDpoly uses one-hot encoded molecule-class features, provided through RoseTTAfold’s 1D track, to guide the generation of appropriate backbone geometry for each polymer type – producing chemically accurate RNA and DNA without compromising protein structure quality. With RFDpoly trained to denoise these different biopolymers, we can specify which polymer class is generated in each contiguous chain during inference (**Fig. 1c**).

**Figure 1.**
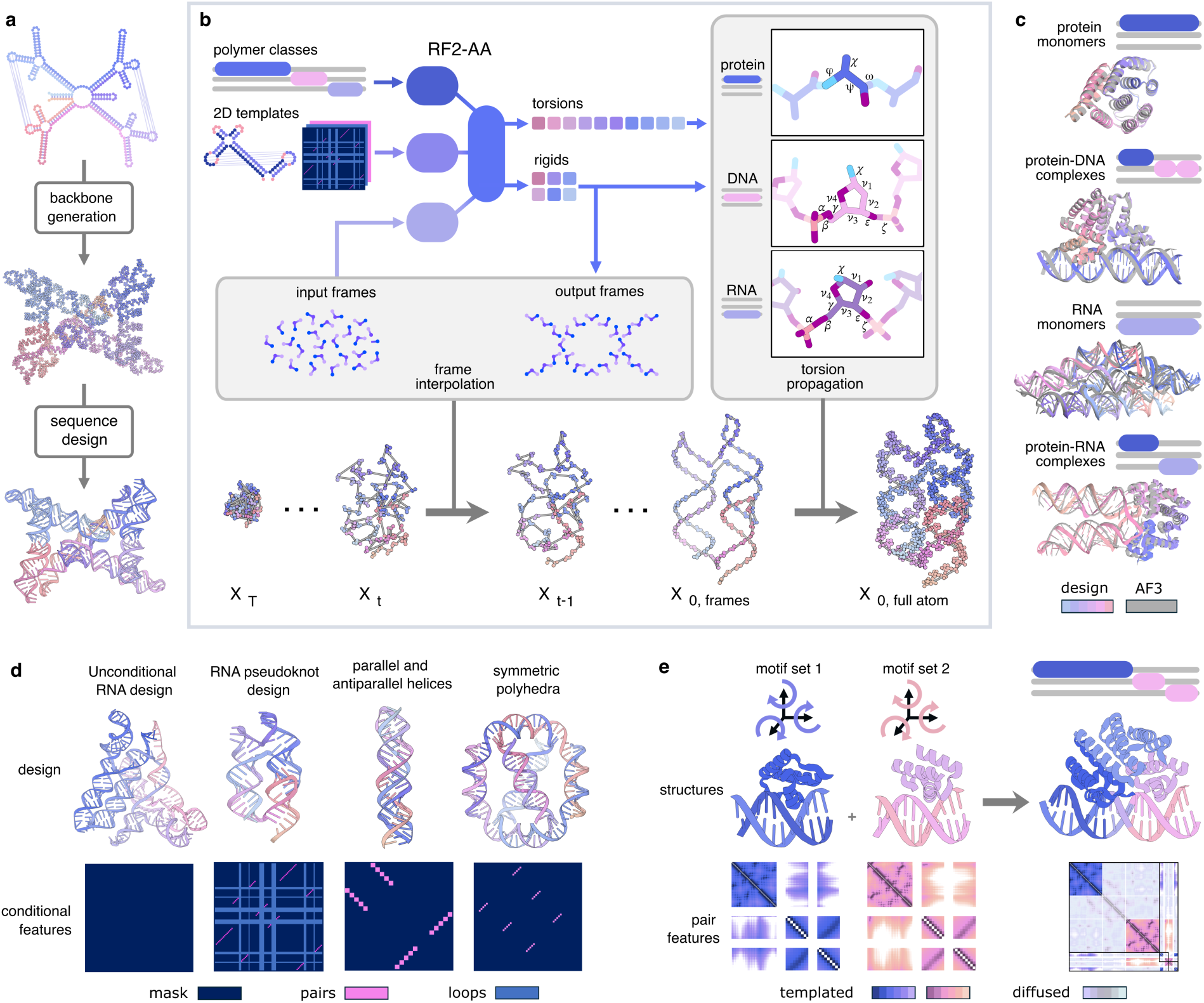
Generalized biopolymer structure generation with RFDpoly. **a**, Standard *de novo* structure design pipeline, consisting of backbone generation followed by sequence design. **b**, RFDpoly denoises backbone frames using RoseTTAfold’s 3D track, while conditional information from the 1D and 2D tracks (such as polymer class and base-pair patterns, respectively) guides the generation toward desired structures. Full sugar–phosphate backbone coordinates are constructed by propagating torsion angles about frames, using connectivity graphs defined by sequence predictions. **c**, Polymer-class labels provided at inference instruct the model to generate chemically accurate structures appropriate for each molecule type. *In silico* predictions with AlphaFold3 show high self-consistency for proteins, DNA, and RNA. **d**, Structural control via base-pair templating enables RFDpoly to generate nucleic acid structures with diverse topologies. User-specified controls define templates for RoseTTAfold’s 2D track. **e**, Hierarchical design in RFDpoly uses 2D templates for motif structure, allowing global placement to be inferred during the denoising process.

To enable RFDpoly to condition on base-pairing networks when generating 3D RNA structures, we expanded RoseTTAfold to accept base pair partner labels through its 2D-track (**Fig. 1b**). These conditional features allow RFDpoly to generate backbones that possess specified secondary structure patterns. To enable user guidance over structure generation in ways that are relevant to nucleic acid design, we provide several routes for controlling secondary structure features at inference time **(Fig. 1d)**. Secondary structure strings (in the form of dot bracket notation) can be provided to generate structures with specific pseudoknot topologies, allowing users to generate 3D models for RNA puzzles^3,25,26^ or other sources of secondary structure templates. Beyond generating canonical single-partner base pairs, users can specify whether pairs are parallel or antiparallel, giving precise control over tertiary structure features such as triple helices or G-quadruplexes. Lists of paired sequence regions can be specified, with cyclic- or repeat-symmetry, to reflect design patterns often seen in nucleic acid origami.

We train RFDpoly using a randomized masking scheme in which training examples are divided into *motif* regions (containing ground truth structure information) and *diffused* regions – to emulate design tasks such as motif-bridging, docking, or generating new structure around central motifs. To enable RFDpoly to generate both protein and nucleic acid structures, and assemblies containing both polymer types, we train the model using datasets consisting of proteins, RNA, and DNA either as monomers or as mixed polymer complexes. While previous versions of RFdiffusion only generate structure within protein chains, keeping nucleic acids as fixed motifs, the mask generators for RFDpoly treat all polymer classes equally, denoising noised regions within nucleic acid chains as well as proteins.

Once backbone structures have been generated, sequence design is performed using two methods. In the first approach, NA-MPNN is used to assign sequences to complete diffusion-generated backbones^22^, and PyRosetta is then used to build and repack sidechains. In the second approach, sequence-structure codesign, RoseTTAfold sequence predictions are autoregressively decoded along with rigid-body and torsion parameters to produce full atom sidechain representations. Sequences designed with NA-MPNN were predicted to fold with better self-consistency metrics, but the presence of side chains from codesign trajectories allowed immediate filtering of structures for target base pair pattern satisfaction, without going through the process of MPNN-design and PyRosetta repacking.

Designed sequences were filtered based on similarity between design models and predicted structures at multiple levels. Initial filtering was based on predicted secondary structure self-consistency using RibonanzaNet^27^; similarity between designed and predicted secondary structure was evaluated using F1-scores^28^. Tertiary structures of designs with high F1 scores were predicted using Chai-1^29^ or AlphaFold3^30^, and compared to design models, using GDT^28,31,32^ or TM-score^33^ as metrics for structure similarity. We use the term “designable’’ below to describe designs for which the predicted structure matched the design model.

In unconditional structure generation calculations, the generated structures maintained high designability across increasing lengths, with little drop-off in secondary structure self-consistency metrics, and minimal steric clashing even in large-scale generation tasks (**Extended Data Fig. 1a**). Tertiary structure *diversity* was evaluated by comparing inter- and intra-group distributions of 3-dimensional fold similarities for diffusion-generated structures against native RNAs from structural databases, or against other diffusion-generated structures; tertiary structure *designability* was evaluated by comparing designed RNAs to AF3 predictions of their MPNN-designed sequences. Pairwise alignment of RNA structures (within 20% sequence length) from both generated and native sets (∼1,700 RCSB RNA structures) using USalign^33^, yielded distributions of tertiary structure similarity (given by TM-score) across various length scales (**Extended Data Fig. 1b**). While the generated structures of short RNAs had folds resembling those in the PDB (e.g., tRNAs), diversity increased rapidly for structures longer than 120 bases. RFDpoly-generated structures were dissimilar from native RNAs, while still having high AF3-predicted self-consistency (**Extended Data Fig. 1c**), indicating that the model can explore novel folds on the global level, without loss in designability.

### Experimental evaluation in the Eterna OpenKnot challenges

To benchmark the ability of our method to generate structures with specific base-pair patterns, we participated in the Eterna OpenKnot challenges^26^. Base-pairing templates from a set of 57 pseudoknot puzzles were used to condition denoising trajectories **(Fig. 2a)**, and sequences were assigned using NA-MPNN. For all puzzles attempted, reference wild-type sequences were provided for comparison, and many puzzles had experimentally determined 3D structures, which were also redesigned with NA-MPNN. To experimentally characterize the secondary structures of designed sequences, the OpenKnot organizers used SHAPE-seq, which generates reactivity profiles that distinguish between base pairing and non-base pairing regions^27, 34,35,36^.

**Figure 2.**
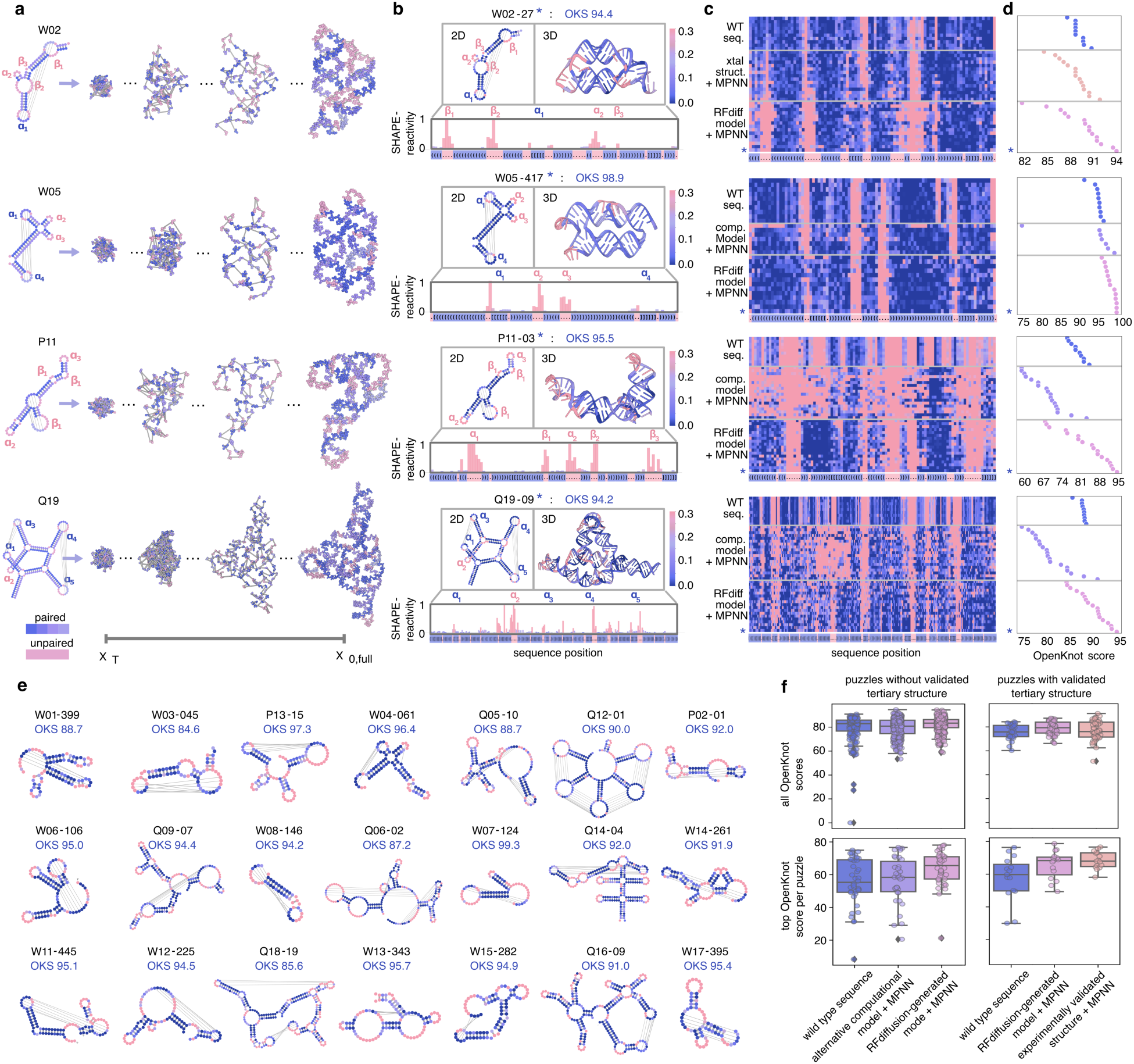
Experimental validation with SHAPE-seq. **a**, Denoising trajectories for representative RNA puzzles, where puzzle templates specify base-pair patterns and unpaired loop regions. **b**, SHAPE profiles for selected designs show high reactivity in unpaired regions (structures colored by reactivity), consistent with target secondary structure templates. Annotated structural features include hairpins (α) and bulges (β). **c**, Experimental SHAPE profiles compiled for wild-type sequences, MPNN-redesigns of alternative backbones (experimental or computational), and MPNN-designs of diffusion outputs. Selected designs (*) best matched target structures. **d**, Target OpenKnot scores derived from SHAPE profiles. **e**, SHAPE profiles mapped onto puzzle secondary structures with corresponding OpenKnot scores, for selected puzzles. **f**, Distribution of all Target OpenKnot scores (top) and best scores achieved per puzzle (bottom), separated by puzzles with experimentally determined 3D structures (left) and those with only computational models (right).

For clarity, we first describe the design approach and the comparison to experimental data in more depth for a specific case, puzzle Q19 (SV_r7_240_4), which consists of five hairpins (α_1_– α_5_). In the target structure, hairpin pairs (α_1_ : α_3_) and (α_4_ : α_5_) form kissing-loop interactions, while hairpin α_2_ remains unpaired (**Fig. 2a**, left). Starting from random noise, the base pair-conditioned diffusion denoising trajectory progressively builds an RNA structure adhering closely to these predefined contacts and hairpin geometries (**Fig. 2a**, right). Hosts of the OpenKnot competition experimentally assessed this design using selective 2’-hydroxyl acylation analyzed by primer extension sequencing (SHAPE-seq), which reports on nucleotide flexibility and accessibility by chemically modifying unpaired or flexible nucleotides. Modified nucleotides impede reverse transcription, allowing sequencing readouts to quantify site-specific reactivities.

Regions displaying high SHAPE reactivity correspond to unpaired or loop nucleotides, whereas paired bases are protected from modification and thus exhibit low reactivity. To show how reactivity data map back to structural features, we color secondary and tertiary structures by reactivity profile for a selected design (*) (**Fig. 2b**). Consistent with expectation, we observe low reactivity in the paired kissing loops, but high reactivity in the unpaired stem loop bases (α_2_).

Thus, the SHAPE-seq reactivity profiles indicate that the design secondary structure closely matches the target secondary structure (horizontal axes, with desired paired and unpaired positions colored in blue and red, respectively). To systematically evaluate performance across multiple design strategies, SHAPE-reactivity profiles were compiled into two-dimensional heatmaps (**Fig. 2c**), comparing our diffusion-based method with alternative backbone redesigns and wild-type RNA sequences. The selected design (*) displayed optimal agreement with the target structure, as quantitatively captured by the highest achieved target OpenKnot score among designs for puzzle Q19 (**Fig. 2d**).

Overall, SHAPE-seq experimental reactivity profiles for targeted puzzles were close to the input base pairing profiles **(Fig. 2c)**. *Target OpenKnot Scores*^37^ were used as an aggregate metric to summarize how well each design folded into the target secondary structure **(Fig. 2d-e).** For most puzzles attempted, at least one sequence from MPNN-design of RFDpoly-backbones outperformed the wild-type sequences and MPNN-redesign of experimental reference structures (**Fig. 2f**, **Extended Data Fig. 2a**-c).

These results show that RFDpoly can generate 3D models across a wide range of target secondary structures. User-input enables the generation of designable backbones for a diverse set of pseudoknot topologies without loss in designability (**Fig. 2e**). The diversity and complexity of secondary structures designed indicate that RFDpoly can explore a complex fold space beyond that which is accessible to origami approaches or even observed in native systems.

### Design of pseudocycles

To further assess RFDpoly’s design capabilities, we sought to create larger novel RNA architectures. We focused on designing *RNA pseudocycles*: large, monomeric RNA structures composed of nearly cyclically repeating subunits connected in a single continuous chain. These folds were chosen as design targets because their large size, relative to our previously designed RNA pseudoknots, would enable characterization via electron microscopy, and their nearly symmetric morphologies would be easily identifiable during screening. We chose single-chain pseudocycles rather than cyclic assemblies to avoid the complications of multimeric assembly, enabling direct application of *in silico* screening metrics optimized for single-chain RNAs.

To generate large pseudocycles with numerous intra-chain contacts to reduce structure flexibility, we developed a secondary structure template generation protocol that accepts graph-based definitions of regional pairing and efficiently searches for fitting base-pair patterns, diversified using random sampling of subregion lengths (**Fig. 3a**). Once candidate base pair templates were generated, they were provided to RFDpoly as 2D templates to guide the denoising process. Pseudocyclic RNA backbones were generated using symmetric noise propagation during denoising, while maintaining backbone connectivity between subunits in encoded bond features. After backbone generation and sequence design, *in silico* predictions of tertiary structure with AlphaFold3 showed conformational variability between models, while having matching secondary structures between designed and predicted structures (**Fig. 3b–c**). Synthetic genes were obtained and *in vitro*-transcribed for eight 372-base designs, all sharing a common twofold pseudocyclic symmetry separated by hinge-like loops.

**Figure 3.**
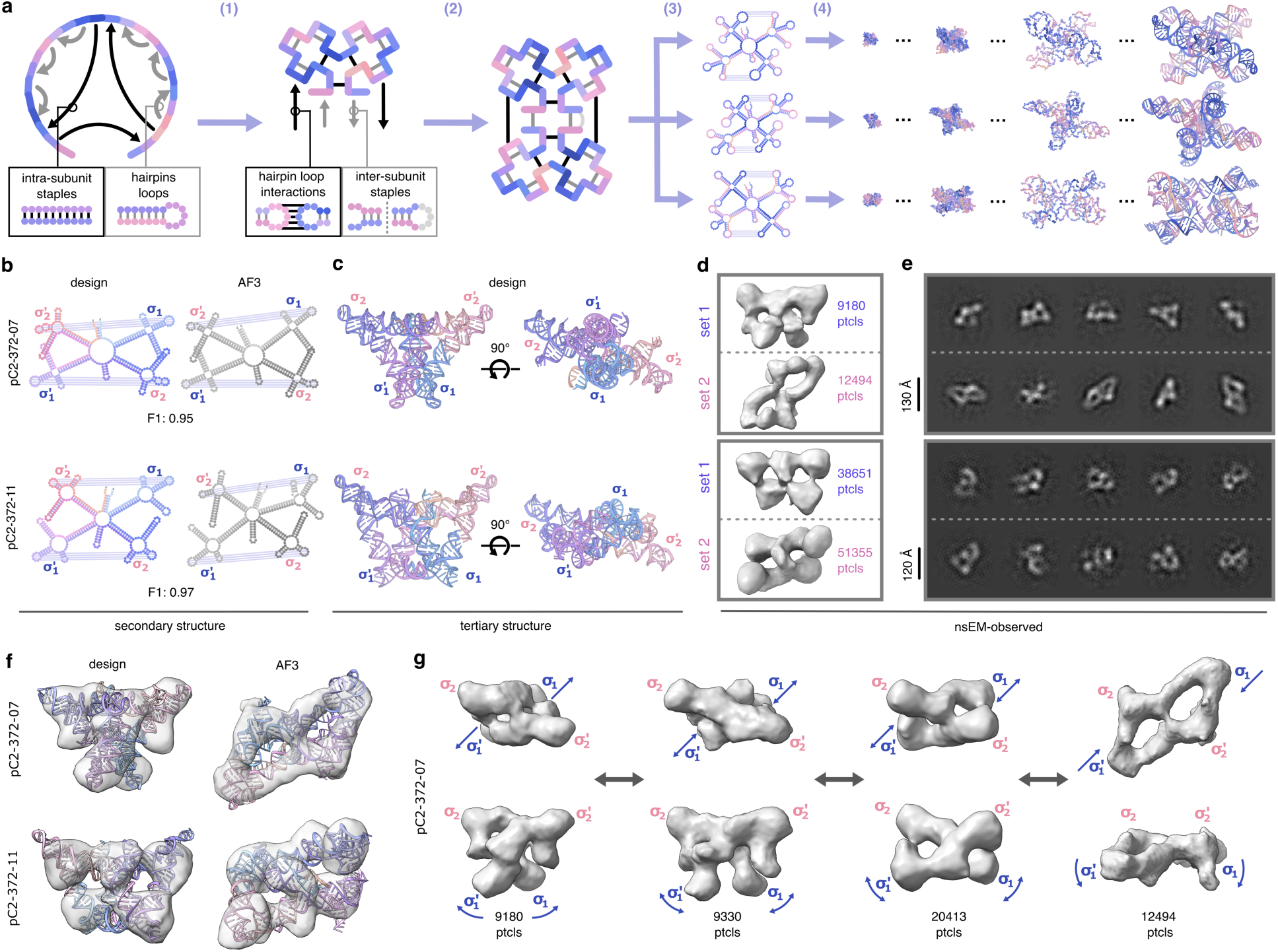
Design and characterization of 372-base pseudocycles. **a**, Workflow for generating base pair templates and RNA pseudocycles: (1) initialize subunit segments and assign *intra*-subunit partner regions; (2) propagate subunits symmetrically and assign *inter*-subunit pairs; (3) randomly vary segment lengths to diversify structures; (4) run RFDpoly denoising with secondary structure conditioning. **b**, Comparison of secondary structures and F1-similarity scores between design models and AlphaFold3 predictions for selected designs (top: pC2-372-07, bottom: pC2-372-11). **c**, Tertiary structures of selected designs. **d**, 2D class averages from nsEM data. **e**, 3D reconstructed nsEM volumes showing butterfly-like folds consistent with design models and AF3 models. **f**, Design models and AF3 models fit into reconstructed volumes. **g**, Conformational variability in pC2-372-07 visualized across multiple nsEM volumes.

Characterization with negative stain electron microscopy (nsEM) revealed well-formed C2 pseudosymmetric structures for four designs (**Extended Data Fig. 3a**-c). Observed morphologies were polydisperse, with the butterfly-like folds sampling multiple conformational states, and the two rigid side-domains folding about their central hinge loops. For two of the four designs (pC2-372-07, pC2-372-11), class averages for one of the primary observed particles (denoted *set-1*) matched the design model, while others (*set-2*) more closely matched the AlphaFold3 predicted models (**Fig. 3d–f**). For the remaining designs, reconstructed volumes did not closely resemble either the design or predicted structures, while nsEM revealed alternative pseudosymmetric folds (**Extended Data Fig. 3d**). All of the nsEM-characterized designs had similar secondary structure features (defined by four characteristic cloverleaf folds: σ_1_, σ_2_, σ_1_’, σ_2_’), and for each design with nsEM 3D reconstructions, one of the primary observed states had the target double-triangle fold (*set-1*). Fitting of design models into 3D-reconstructed volumes for particle *sets-1* demonstrated structural consistency within this subpopulation (**Fig. 3f**). To further characterize the hinge-motion between the open and closed conformations, we classified particles into multiple 3D volumes, which revealed connectivities between the designed conformation and alternative states (**Fig. 3g**).

We next designed smaller (240 nt) compact RNA pseudocycles, aiming for well-defined tertiary structures by enriching for supportive base pair contacts and iteratively refining the overall structure by recycling the most well-ordered regions of the models back into RFDpoly as base pair templates, over multiple rounds of design and selection. These shorter-length sequences could be characterized with SHAPE-seq, allowing us to compare experimental results between chemical probing and electron microscopy. Fifty designs with two- or three-fold pseudocyclic symmetry were selected for SHAPE-seq characterization based on high *in silico* self-consistency metrics. Overall, there was strong agreement between experimental reactivity profiles and target secondary structures (**Fig 4a**), with 42 designs achieving *Target OpenKnot Scores* above a target threshold of 0.8 (**Extended Data Fig. 4a**). Of these, 19 were selected for further characterization by electron microscopy, based on whether Chai-1^29^ or AlphaFold3^30^ predictions resembled the design models. RNA transcribed from DNA templates was screened by nsEM, and the most structured candidates were further characterized by cryo-EM. Three designs (pC2-240-02, pC2-240-03, pC2-240-04) had good SHAPE-reactivity profiles (**Extended Data Fig. 4b**) and well-defined 2D classes from cryo-EM (**Fig. 4b**, top; **Extended Data Fig. 4c-d**).

**Figure 4.**
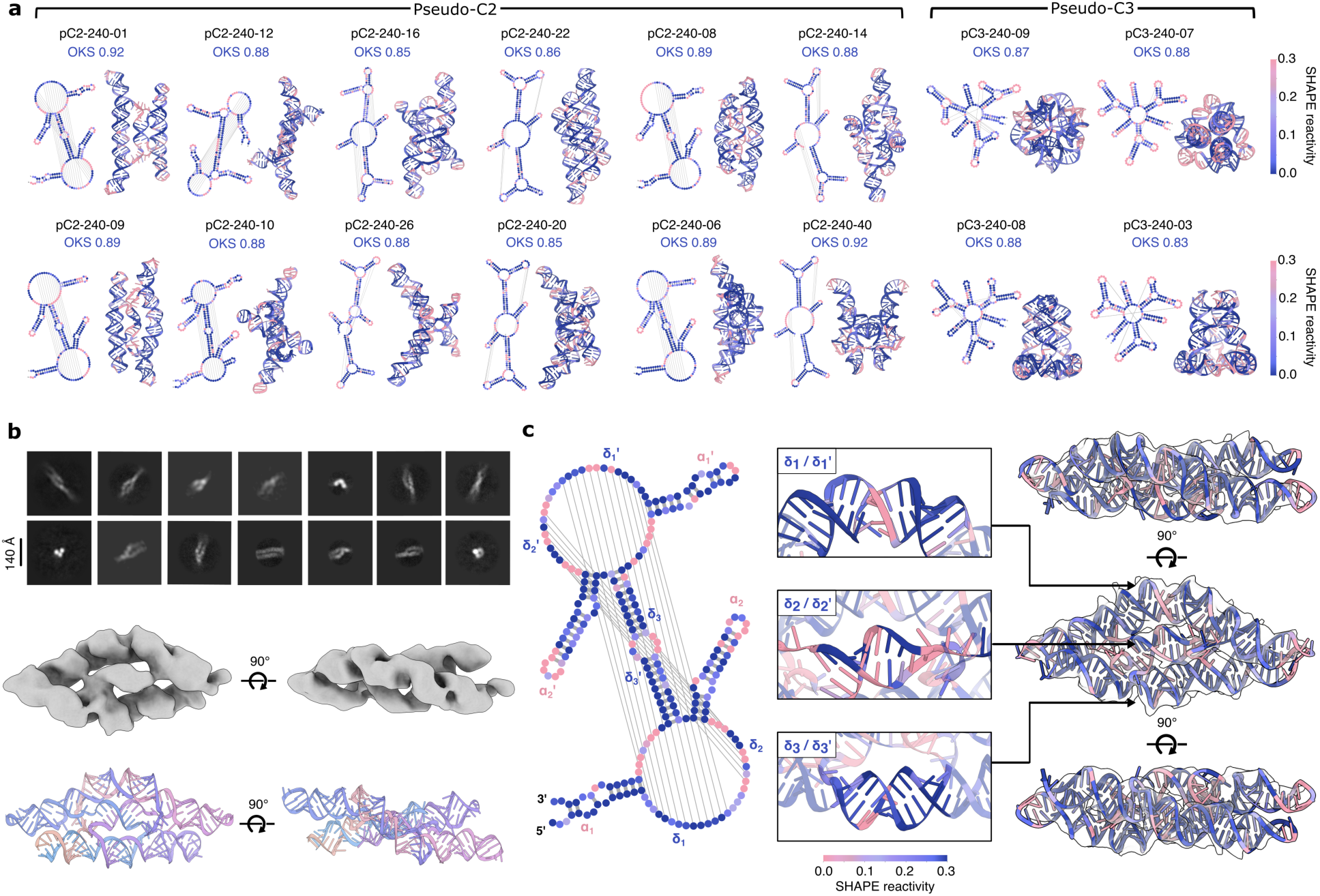
Characterization of small, compact pseudocycles. **a,** Representative compact pseudocycle designs; secondary (left) and tertiary (right) structures colored by SHAPE-seq reactivity. **b**, Cryo-EM 2D classes (top), and 3D reconstructed volume (middle) of designed pseudocycle, pC2-240-04, revealing the overall fold of the design model (bottom). **c**, SHAPE reactivity profile of pC2-240-04, mapped onto the secondary structure (left) and tertiary structure (right), fit into the cryo-EM volume. Three central helical domains (δ_1_, δ_2_, δ_3_) are highlighted (middle), showing crossover base pairs in the secondary structure and corresponding strand interweaving in the tertiary fold.

For pC2-240-04, cryo-EM characterization led to a 6.6Å reconstructed volume, with clearly observable helical pitch and a global fold matching the design model (**Fig. 4b**, middle and bottom). The experimental SHAPE-reactivity profile matches the designed secondary structure (**Fig. 4c**, left), and the design model could be fit into the cryoEM reconstructed volume (**Fig. 4c**, right). The locations of paired and loop regions observed in the cryo-EM reconstruction are consistent with the SHAPE-reactivity profile. Of particular interest are three central helical domains, δ_1_, δ_2_ and δ_3_ (**Fig. 4c**, middle), with crossover regions in the secondary structure that make this fold a complex high-order pseudoknot (since these central crossover domains can occupy separate locations in 2D secondary structure diagrams, we label paired strands as δ*_i_* and δ*_i_*‘, denoting the upstream and downstream components, respectively). The interweaving of these three helical domains between protruding helical hairpins in the design model is confirmed in the 3D tertiary structure. To assess the novelty of this fold, we aligned the experimentally refined model against all representative RNAs from structural databases (see Methods) and found no close matches: the best hit was to a small region of the plant mitochondrial ribosome (PDB 6XYW, chain A) with a TM-score of 0.329 at 0.55 coverage. The combined validation using SHAPE-seq and cryoEM shows that RFDpoly can accurately create intricate new RNA tertiary structures.

### Motif scaffolding and hierarchical design

We next set out to use RFDpoly to design protein–nucleic acid assemblies. We began by incorporating cognate protein and DNA motif fragments from previous work^38^ into larger assemblies; simultaneously fusing interacting protein (C-to-N) and DNA (3′-to-5′) motifs with newly generated rigid connective structures. We used 2D motif-template conditioning to embed three sequence-orthogonal DNA-binding proteins (DBPs) and their respective target DNAs (**Fig. 5a**) into larger connected assemblies, allowing their global placement and the connections between them to be inferred during the denoising process (**Fig. 5b)**. Base pair templates that specified DNA strand exchange were used to condition the denoising process, and control long-range connectivity (**Fig. 5c,d**), enabling the creation of intricate nucleoprotein complexes with cooperative, multi-contact binding surfaces that would be challenging to achieve with purely parametric approaches.

**Figure 5.**
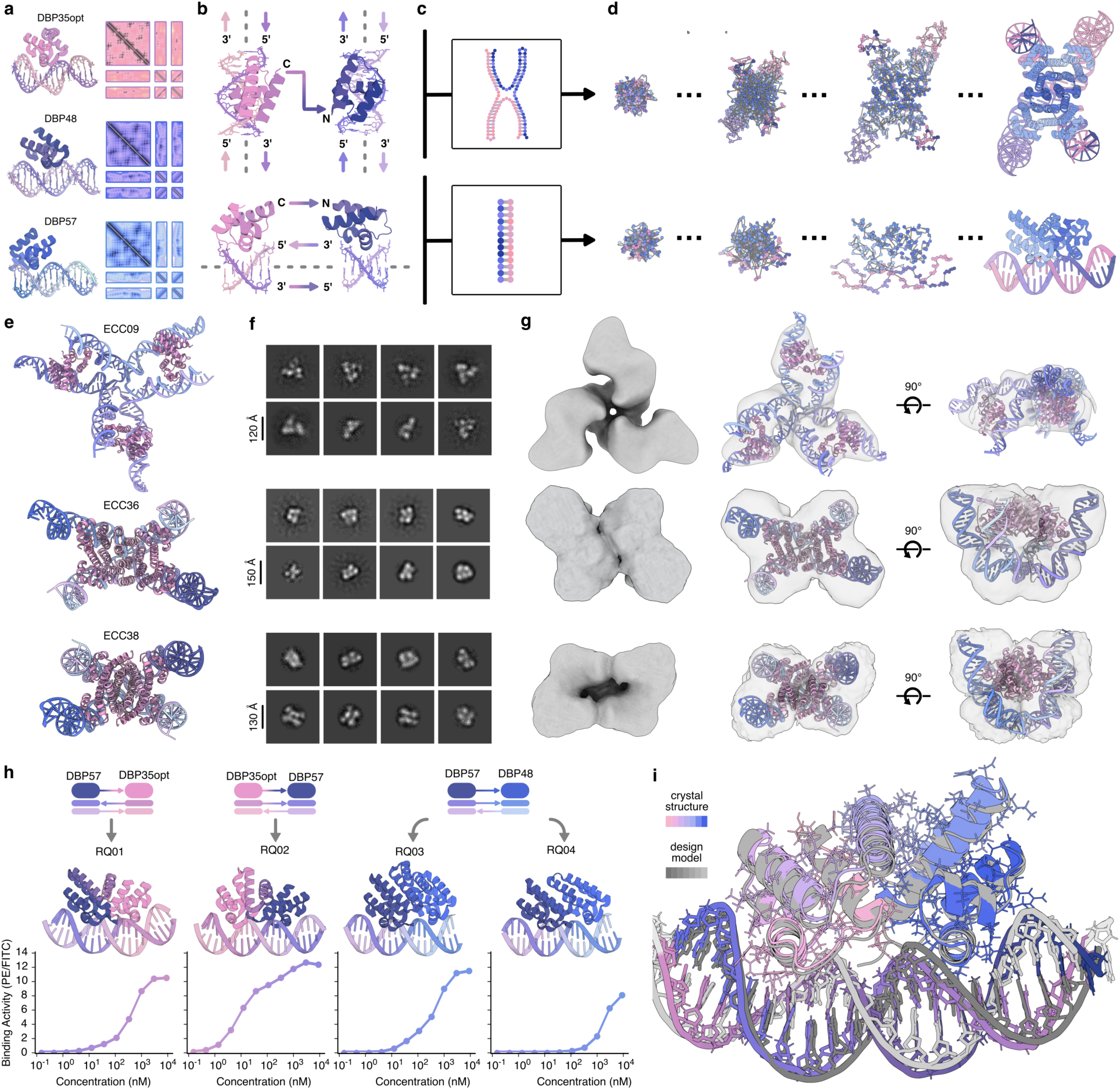
Multi-polymer fusions and motif scaffolding. **a,** DNA-binding proteins (DBPs) and their 2D templates, provided to RFDpoly for motif scaffolding. **b,** Multi-chain protein-DNA fusions: either *inter*-helical fusions bridging two DBPs across different helices (top) or *intra*-helical fusions parallel to the DNA axis (bottom). **c,** Base-pair pattern specification enables topological control over generated DNA interfaces. **d,** Denoising trajectories run with or without symmetry (top and bottom, respectively) to generate complex assemblies. **e,** Design models of three nucleoprotein junctions, with protein domains bound to strand-exchanging DNA hetero-oligomers. **f,** nsEM 2D class averages of selected designs. **g,** nsEM 3D reconstructed volumes shown alone (left), or superimposed over design models (middle and right); final maps refined using symmetry constraints. **h,** Intra-helical fusion of different DBPs in various binding orientations (top) generated larger sequence-specific DNA-binding proteins (middle), which bind to longer stretches of target DNA, as evaluated by yeast-display flow cytometry assays (bottom). Binding quantified as phycoerythrin/FITC ratios. **i,** Crystal structure of DBP-fusion RQ01 superimposed on the design model using USalign^33^ (RMSD = 1.76, TM-score = 0.90).

We first applied this hierarchical strategy to design protein–DNA complexes in which protein homodimers bind to strand-exchanging DNA heterodimers to form hybrid complexes reminiscent of classical Holliday junctions^39^ (**Fig. 5e**). These designs were created using *inter-helical fusions* (**Fig. 5b, top**) which connect two DBPs across different bound helices, acting as rigid “staples” that lock the relative orientation of the DNA helices, reflecting prior strategies in protein-DNA hybrid nanostructures^40^. An initial set of designs was generated in which DBP fusions form bridges across DNA helical extensions; upon characterization by nsEM, these designs showed high conformational variability, likely due to limited buttressing interactions, which complicated structural characterization. Despite uncertainty in the geometries of these complexes, we did observe on-target trimer assembly for one design, ECC09 (**Fig. 5e-g**, top row). Difficulties in characterizing the flexible complexes motivated a second design campaign, in which the generated protein structure not only connects DBPs but also forms a homooligomeric interface with the other protein chains in the complex, to create a more rigid protein-DNA assembly. This second round of protein-protein interface-forming complexes was expressed and again characterized by nsEM, revealing successful assembly for two designs, ECC36 and ECC38 (**Fig. 5e-g**, bottom two rows). The pseudo-symmetric structure of these complexes is shown clearly in 2D class averages (**Fig. 5f**) and 3D reconstructed volumes (**Fig. 5g**), consistent with the bent configuration of the DNA in these strand-exchanging complexes. Thus, RFDpoly can carry out motif scaffolding while simultaneously generating designs with complex secondary structures, providing a route to bridge the worlds of protein design and DNA origami.

We next used the same flexible scaffolding approach to generate new proteins predicted to bind extended contiguous DNA sequences (**Fig. 5h**). This second hierarchical design campaign utilizes *intra-helix DNA fusions* (**Fig. 5b**, bottom) in which DBPs are concatenated along a shared helical axis to create a longer sequence-specific recognition domain, expanding on existing techniques for the modular assembly of DNA-binding proteins^41,42^. By sampling the geometry of the connecting DNA during DBP fusion, the designs can bind DNA in a wide range of conformations (e.g., **Fig. 5i**), in contrast to a previous approach limited to linear, parametrically generated B-form DNA^38^. The newly designed intra-helix DBP fusions were evaluated by yeast-display flow cytometry experiments (**Fig. 5h**, bottom) in which biotinylated dsDNA targets were titrated to generate binding curves, yielding apparent K_D_ values from ∼2 µM (RQ04) to ∼10 nM (RQ02).

We assessed model accuracy by X-ray crystallography, obtaining a 3.0 Å resolution crystal structure of RQ01 bound to its DNA target (**Fig. 5i**). The crystal structure closely matches the design model over the full assembly, including the rigid linkers; notably, the bent DNA conforms to the design, indicating that RFDpoly can accurately design non-ideal nucleic-acid geometries. Together, these results showcase the utility of RFDpoly for generating new DNA-binding proteins specific for extended DNA sequences and the ability to scaffold multiple biopolymer types simultaneously. Overall, our hierarchical design approach allows for the reuse of previously characterized structures to accelerate further design efforts and enables the construction of increasingly complex classes of biomolecular architectures.

## Discussion

The *de novo* design of nucleic acid tertiary structures has been a longstanding challenge due to their intrinsic flexibility and the relatively small number of structures available for training deep learning models compared to proteins. Previous efforts, including adaptations of diffusion models for RNA, have been restricted to short sequences, exhibit limited structural diversity, and lack robust motif scaffolding or experimental validation, leaving the physical plausibility of their outputs uncertain^17,18,19,20,21^. RFDpoly enables user-guided generation of structured RNA with intricate, native-like folds, going beyond the regular origami-style topologies that have long defined the field of three-dimensional nucleic acid design. The use of randomly-sampled motif masks and conditional feature labels during training allows the model to learn local features of molecular structure, and to generalize these globally at inference time. Explicit hydrogen bond conditioning during RFDpoly’s denoising process enables control over the topologies of the structures generated through the installation and enrichment of base-interaction networks. The successful design and cryo-EM validation of a 240-nucleotide RNA pseudocycle demonstrates that RFDpoly can produce novel RNAs with intricate folds. Despite being trained on a relatively small dataset of experimentally determined RNA structures, RFDpoly generalizes beyond known motifs (**Extended Data Fig. 1a**), in some cases producing designs more stable than their natural counterparts (**Extended Data Fig. 2**). The ability of RFDpoly to generate both base pair-conditioned and unconditional structures with high accuracy and diversity suggests that it does not simply memorize motifs from the training data, but instead learns fundamental assembly principles, enabling the exploration and creation of physically viable, previously unobserved RNA folds.

The ability of RFDpoly to generate protein-nucleic acid assemblies enables the design of increasingly complex structures by reusing existing components. Combining the capabilities of RFDpoly to carry out polymer-class conditioning, 2D-templated scaffolding of preexisting motifs, and base-pair templating to guide nucleic acid secondary structures enabled the design of the protein-DNA assemblies in **Fig. 5**. The close agreement of the crystal structure of a designed protein-DNA assembly, RQ01, with the design model showcases the ability of RFDpoly to generate atomically accurate designs and to stabilize irregular (bent) DNA backbone structures. The relative placements of the input motif sets were not pre-defined within a global coordinate frame as in previous work: instead, the model inferred a physically plausible arrangement while generating the connecting backbones, eliminating the need for manual motif positioning, and enabling joint optimization of motif placement and backbone generation.

Despite the many new design capabilities described herein, there are still many avenues to explore to expand our ability to design RNA and nucleoprotein complexes. Developing compact, partner-specific protein-RNA interfaces would unlock fully RNA-centric scaffolds for hierarchical assembly, complementing the DBP-DNA fusions shown in this work. The development of more reliable tertiary structure prediction methods for protein-nucleic acid assemblies would considerably improve *in silico* screening of generated designs^43^. Finally, incorporating greater chemical diversity (small-molecule ligands, noncanonical nucleotides, and post-synthetic modifications) would expand RFDpoly’s generative capabilities and enable the design of programmable sensors and control elements akin to riboswitches. Ultimately, the continued development of generative frameworks like RFDpoly will likely transform the design of structured nucleic acids and proteins, unite protein design and DNA origami methods, and enable rapid, precise engineering of biomolecular assemblies for synthetic biology, therapeutic control, and beyond.

**Table 1:** Crystal structure data collection and refinement statistics.

|  | Design: RQ01 |
| --- | --- |
| Wavelength |  |
| Resolution range | 43.63 - 2.116 (2.25 - 2.12) |
| Space group | C 1 2 1 |
| Unit cell | 113.148 33.745 104.306 90 107.23 90 |
| Total reflections | 115915 (5311) |
| Unique reflections | 36743 (2847) |
| Multiplicity | 3.2 (1.9) |
| Completeness (%) | 82.33 (20.91) |
| Mean I/sigma(I) | 12.02 (0.68) |
| Wilson B-factor | 34.95 |
| R-merge | 0.04013 (1.089) |
| R-meas | 0.04781 (1.473) |
| R-pim | 0.02569 (0.9807) |
| CC1/2 | 0.999 (0.432) |
| CC* | 1 (0.777) |
| Reflections used in refinement | 18082 (751) |
| Reflections used for R-free | 892 (28) |
| R-work | 0.2192 (0.2972) |
| R-free | 0.2473 (0.2885) |
| Number of non-hydrogen atoms | 2418 |
| macromolecules | 2274 |
| ligands | 0 |
| solvent | 144 |
| Protein residues | 166 |
| RMS(bonds) | 0.004 |
| RMS(angles) | 0.55 |
| Ramachandran favored (%) | 98.15 |
| Ramachandran allowed (%) | 1.85 |
| Ramachandran outliers (%) | 0 |
| Rotamer outliers (%) | 1.55 |
| Clashscore | 3.14 |
| Average B-factor | 48.86 |
| macromolecules | 39.69 |
| solvent | 45.94 |

## Acknowledgements

We thank the Eterna project and the Das lab (Stanford, HHMI) for coordination and experimental evaluation of the Eterna OpenKnot challenge. The authors would like to thank Lance Stewart and Lynda Stuart for their work in maintaining operations at the Institute for Protein Design; Luki Goldschmidt and Kandise VanWormer and for maintaining the computational and wet lab resources that were used in this research; Madison Kennedy and Bulat Faezov for technical help in preparation of this manuscript; Han Altae-Tran, Pooja D Bandawane, Andrew C. Hunt, and Florian Praetorius for experimental advice and insightful discussions over the course of this project.

## Author contributions

A.H.F. conceived the study, wrote the manuscript, trained RFDpoly, and developed the RNA design pipeline. A.H.F., R.Q., and E.C. generated experimentally characterized designs. R.Q., E.C., E.S., Y.P., A.C., and Y.H. purified proteins. A.H.F purified RNA. A.J.K., L.M., P.T.K., and D.J. contributed additional code. C.G., A.J.B., P.S.K., R.P., F.D., and D.B. offered supervision throughout the project. B.L.S. and M.A.E. characterized crystal structures. C.W., A.J.B., K.C., A.H.F., A.C., and Y.H. performed electron microscopy and data processing. A.J.K. developed and trained NA-MPNN. L.M. and F.D. trained RF-NA.

## Extended Data and Figures

**Extended Data Figure 1:**
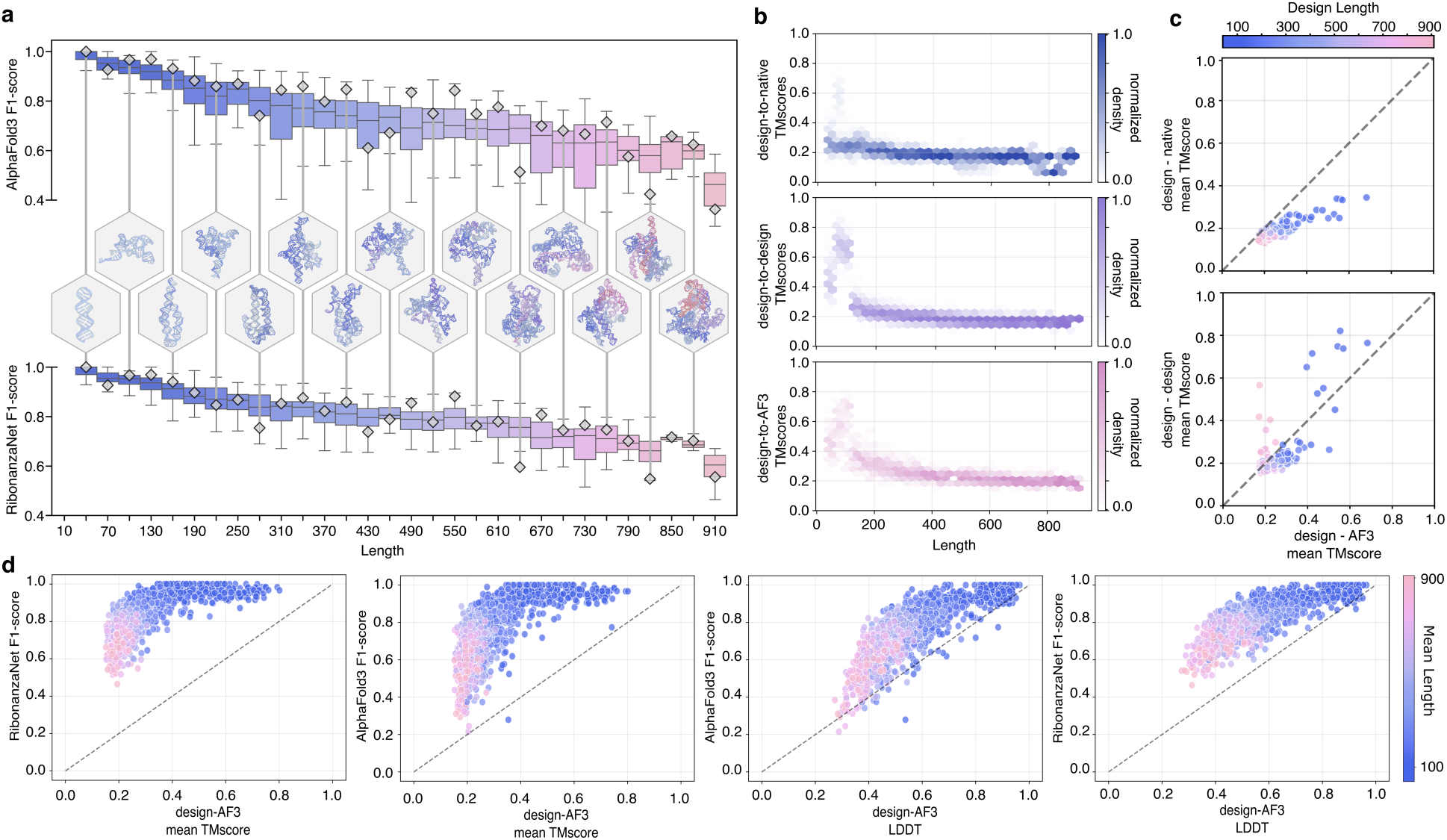
*In silico* performance of unconditional RNA structure generation. **a**, Predicted secondary structure accuracy as a function of sequence length. Top: F1-score similarity between AlphaFold3 predictions and design model secondary structures. Bottom: F1-score similarity between RibonanzaNet predictions and design model secondary structures. Representative designs are shown in hexagons, with vertical connectors to their corresponding F1-score markers. **b.** Distributions of pairwise TM-scores across different length ranges (coverage threshold 0.8). Top: RFDpoly-generated design models aligned to native RNAs. Middle: designed structures aligned to other designed RNAs. Bottom: designed structures aligned to AF3-predicted RNAs. **c.** Per-design comparison of TM-scores to show structural similarity between groups vs each design’s AF3-aligned TM-scores as a self-consistency metric (shared x-axes). Top y-axis: TM-scores of designed backbones aligned to similar-length native structures. Bottom y-axis: TM-scores of designed backbones aligned to other designed backbones. **d.** Comparison of *in silico* predicted secondary structure accuracy metrics (y-axes, RibonanzaNet F1-score or AF3 F1-score) with tertiary structure accuracy (x-axes, TM-score or LDDT from alignment to AF3 model).

**Extended Data Figure 2:**
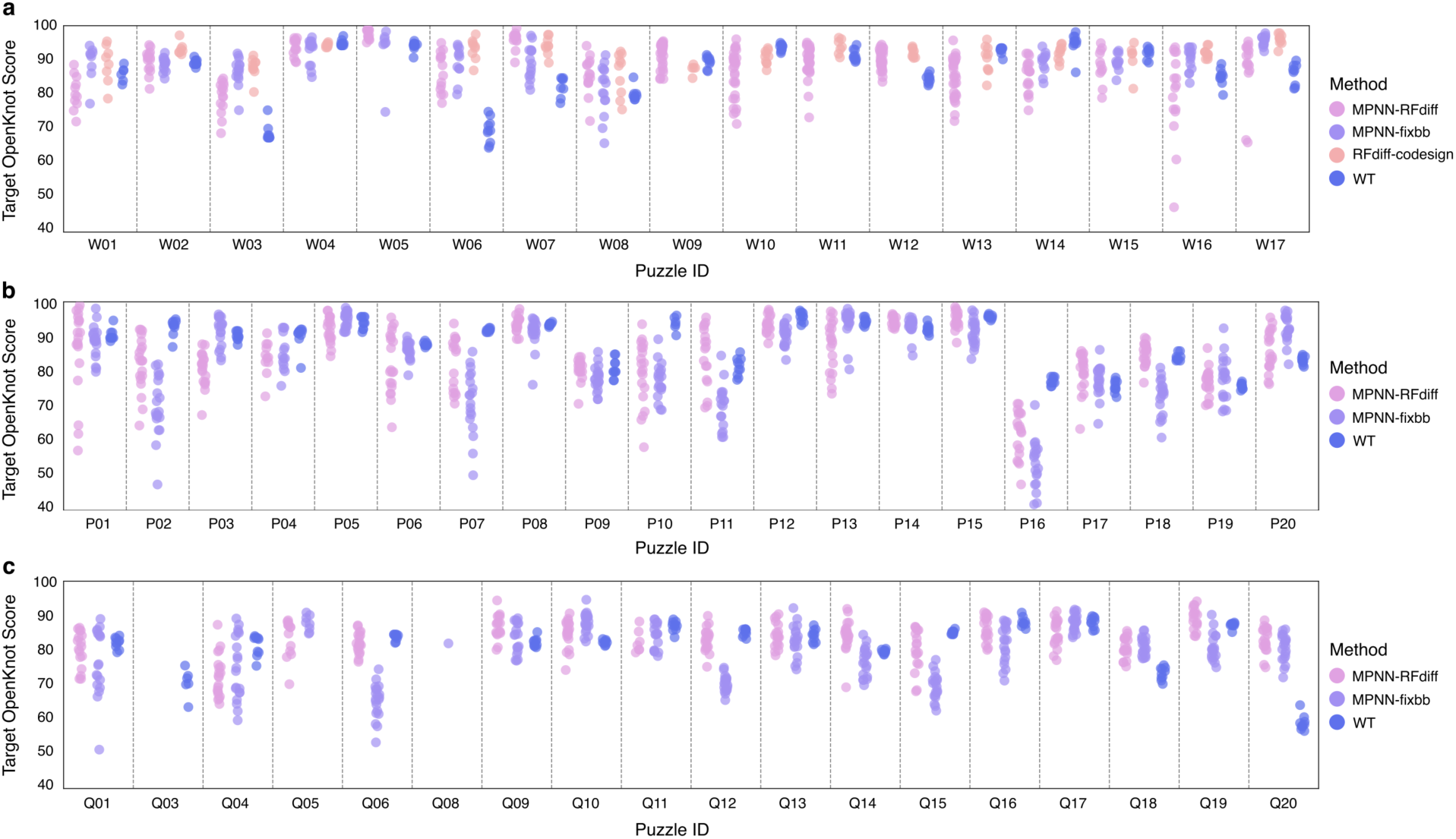
Target OpenKnot scores across competition rounds. **a**, OpenKnot round 6. **b**, OpenKnot round 7a. **c**, OpenKnot round 7b. Note: *MPNN-fixbb* refers to both experimental structures and alternative (non-RFDpoly generated) computational models redesigned by MPNN, depending on the puzzle and what alternative structures were available for each puzzle. *RFdiff-codesign* refers to the autoregressive decoding of RFDpoly’s own sequence predictions over the last 40 steps of 50-step denoising trajectories.

**Extended Data Figure 3:**
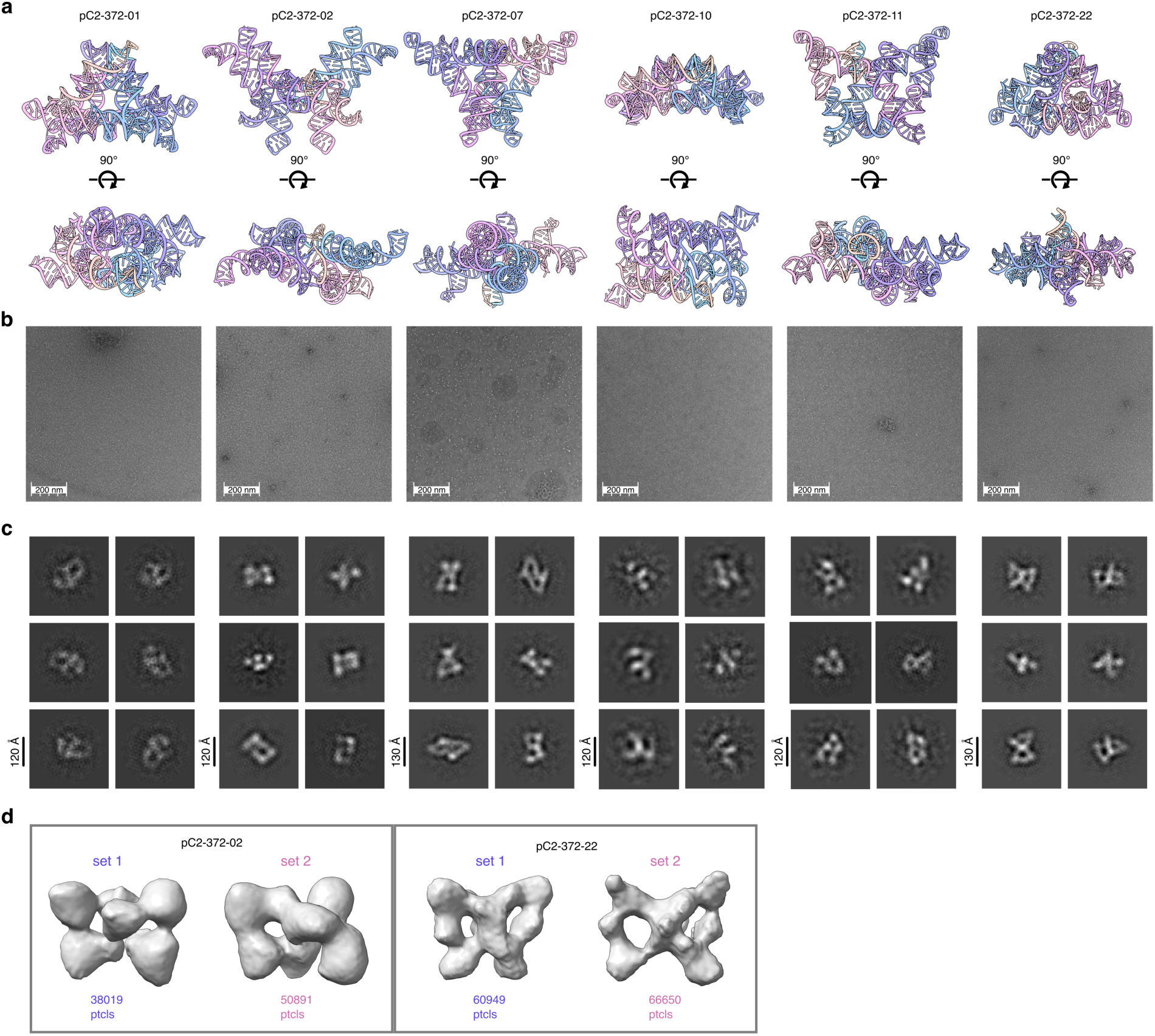
nsEM data for *butterfly fold* (372 nt) pseudocycles. Data for six designs (pC2-372-01, pC2-372-02, pC2-372-07, pC2-372-10, pC2-372-11, pC2-372-22). **a**, Design models shown in side-views and top-down views. **b**, Representative negative stain micrographs. **c**, 2D class averages. **d**, Example 3D reconstructed volumes for classes corresponding to two conformationally-distinct particle sets, shown for pC2-372-02 (left) and pC2-372-22 (right).

**Extended Data Figure 4:**
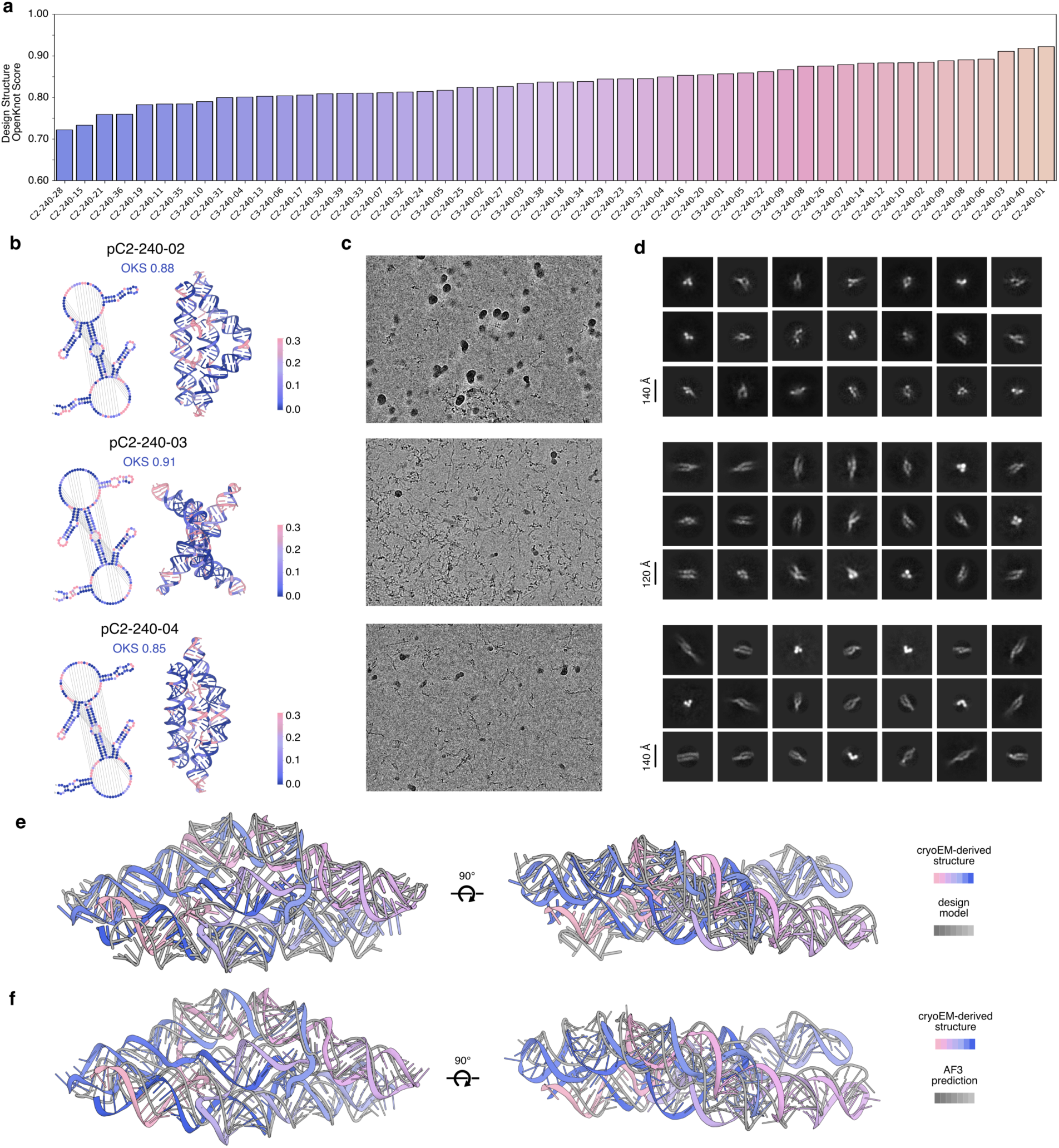
experimental data for *compact* (240 nt) pseudocycles. **a**, Target OpenKnot scores (agreement of SHAPE reactivity with design-model secondary structures) for all 50 compact pseudocycle designs. **b**, Selected designs (pC2-240-02, pC2-240-03, pC2-240-04); secondary and tertiary structures colored by SHAPE-seq reactivity. **c**, Representative cryo-EM micrographs, and **d**, 2D class averages for selected designs. **e,** Comparison of the design model and ISOLDE-refined model, aligned by USalign (RMSD=5.35 Å, TM-score=0.58). **f,** Comparison of the best fitting AlphaFold3 model and ISOLDE-refined model, aligned by USalign (RMSD=5.64, TM-score=0.51).

## Methods

### I. Model Development and Training

#### Motif templating

In RFDpoly, motif geometry is encoded as pairwise template features (2D track) and torsion vectors (1D track), while the 3D track receives only denoised frame coordinates and self-conditioning. This enables motif sets to be grouped and placed independently of a global coordinate frame. Both diffused and motif regions of training structures were subject to random perturbation of frame rotation and translation, since motif geometry was templated exclusively through RoseTTAfold’s pair track using a two-dimensional motif masking scheme, as described previously^44^. Motif chunks were sampled as islands within input structures, with subsets randomly selected to have their relative distances and orientations locked or resampled, which emulated pairwise grouping of motif sets at inference time. While training the model to predict atom placement via torsion propagation was necessary for placing the many nucleic acid backbone atoms in diffused regions, even protein structure generation benefitted from proper torsion predictions in motif regions: our use of pairwise templating necessitated the use of motif torsion vectors to correctly reconstruct full sidechain atom sets within the coordinate frame of the newly generated structure. In previous versions of RFdiffusion, motifs remained fixed in their global frame, allowing sidechain and other atoms to be directly replaced from input structures after diffusion. In our 2D motif templating, motif placements arise during denoising and thus cannot use sidechain replacement. Although backbone frames can be defined from inter-residue distances and orientations^45^, full backbone and sidechain generation via torsion predictions required expanded training tasks, enabling the model to integrate distance maps, torsion templates, and sequence for full-atom structures.

#### Multimodal training

The standard diffusion task is to predict a ground-truth structure from a noised input, with the sequence masked in diffused regions and shown in motif regions. We added additional randomly sampled tasks: (i) predicting full sidechain coordinates from noised structure plus ground-truth sequence and torsions, and (ii) predicting backbone, sequence, and torsions from corrupted backbones to populate all atoms in both diffused and motif regions (the latter in a distinct global reference frame). This multimodal regime enhanced two design functions: (1) respecting full-atom 2D-templated motifs rather than just backbones, and (2) predicting sequences for generated structures, enabling autoregressive sequence-structure codesign during denoising. To handle the added complexity of nucleic acid backbones and 2D-templated motifs, we introduced auxiliary losses (beyond translation and rotation) to score torsion angle predictions and atom placement in local frames.

#### Base-pair feature encoding

For base pair conditioning, we encode a three-channel one-hot tensor, *A* ∈ {0,1}*^L^*^×*L*×3^ on the 2D track: [1,0,0] = *unspecified* (no constraint), [0,1,0] = explicitly *non-pairing*, and [0,0,1] = *paired* for nucleotide pair (*i*, *j*). Tensor *A* is symmetric, the diagonal is set to *unspecified*, and all entries default to *unspecified* unless base pair information is provided, and the 2D encoding permits definition of arbitrary-order pseudoknots. Per-nucleotide encoding was used instead of previously used block-adjacency conventions to indicate regional pairings, because block-level submatrices could not explicitly define the orientation of paired strands without an additional orientation feature. In contrast, orientation can be explicitly defined for each paired region, as whether strands are oriented parallel or antiparallel is reflected by whether their corresponding submatrices are diagonal or the anti-diagonal, respectively. This fine-grained feature encoding provides explicit control over base-pairing arrangements and enables the generation of irregular nucleic acid structures such as triple helices or G-quadruplexes.

#### Inference-time controls

A detailed outline of structural control methods, with many examples, covering multi-polymer design, 2D-templated motif scaffolding, and nucleic acid secondary structure conditioning, will be provided as code documentation and demonstrated through design tutorials, available at the time of publication.

### II. Computational Design Protocols

#### RNA pseudocycle design

Base pair templates matrices were generated from coarse-grain topological specification, as shown in **Figure 3a**. These templates were used to condition RNA structure generation using symmetrically propagated noise and pseudo-symmetric single-chain connectivity, as described previously^46^.

#### Design of linearly-fused protein-DNA complexes

Three *de novo* DNA-binding proteins (DBP35opt, DBP48, DBP57) bound to their target DNAs (TGCACAT, GCCGC, ATCCAGA)^38^ served as scaffolds for multi-chain protein–DNA fusions in parallel orientation (**Fig. 5b, bottom**). 2D motif templating enabled the relative placement of the complexes during denoising. A total of 6,000 diffused designs were generated. DNA linkers were specified at inference, while protein linkers were designed with LigandMPNN^24^. Structures were predicted with RoseTTAFoldNA (RFNA)^16^ and filtered by pLDDT > 0.87, backbone RMSD < 2.2, and protein all-atom RMSD < 3.5 (after alignment via all DNA atoms). Ninety-three designs passed these filters, including RQ01–RQ04 (**Fig. 5h**).

#### Design of semi-symmetric protein-DNA complexes

Semi-symmetric DBP fusions were designed in RFDpoly with symmetric noise, using motif scaffolding to place two DBP motifs per protein chain and their cognate motifs in DNA chains. Secondary structure conditioning was used to give DNA chains their characteristic strand-exchanging topology. An interfacial contact potential promoted protein-protein contacts in DBP fusion regions during denoising. Protein and DNA sequences were designed with NA-MPNN. Symmetric constraints guided protein sequence-design, while asymmetric sequence-design was used for DNA chains in order to produce DNA heterodimers or heterotrimers that would disfavor self-association. Some designs had their sequences further diversified by using ProteinMPNN^23^ to redesign specific residues at protein-protein interfaces. Forty complexes were ordered and tested: 29 with disjoint protein domains and 11 with symmetric protein-protein interfaces.

#### Sequence design and sidechain generation

Sequence design could be performed using two methods. (1) RFDpoly-generated backbones could undergo inverse-folding sequence assignment using NA-MPNN^22^; PyRosetta was then used to thread MPNN sequences onto backbones, build sidechains, and perform quick repacking for reasonable placement. (2) Alternatively, RoseTTAfold’s own sequence predictions could be used to autoregressively assign sequences during the denoising process, combining rigid-body and torsion parameters with sequence identity to produce full-atom models.

### III. Experimental Methods

#### RNA transcription and purification

DNA templates were purchased from Twist and GenScript. In vitro transcription, purification, and refolding followed published protocols^47^. Transcription reactions used reagents from the NEB hi-scribe™ reaction kit. IVT products were purified with the QIAGEN RNeasy kit. Refolding was done in 50 mM Na-HEPES (pH 8.0) by heating to 90 °C for 3 min and cooling for ≥10 min at room temperature. Samples were diluted to a final buffer of 10 mM MgCl₂. All RNAs carried a 5′ GG leader to promote transcription, adding 2 nt to each synthesized monomer.

#### Eterna OpenKnot challenges

Puzzle base pair templates were given by Eterna OpenKnot puzzles in dot-bracket notation. These strings were fed to RFDpoly to produce backbones that could then be designed by NA-MPNN^22^. Sequences were filtered for *in silico* self-consistency using RibonanzaNet secondary structure predictions^27^. The OpenKnot organizers experimentally assessed the secondary structures of designed sequences using SHAPE-seq^34,35,36^.

Additional information about the Eterna OpenKnot challenge (including puzzles and secondary structures) is available at: https://eternagame.org/challenges/11843006

#### Yeast expression of linearly fused DNA-binding proteins

DBP fusion sequences were optimized for *S. cerevisiae*, synthesized as IDT E-blocks with pETCON3 (Addgene #45121) adaptors, and cloned/transformed by in vivo homologous recombination. Specifically, linearized pETCON3 vector was mixed with LiOA (5.5 ng/µL vector, 0.67 M LiOA). 3 µL of pETCON3–LiOA solution and 5 µL of 10 ng/µL E-block inserts were added to 96-well PCR plates and incubated for 5 min at room temperature. 30 µL PEG–LiOA solution (43.3% PEG 3,350, 0.13 M LiOA) and 10 µL competent EBY100 yeast were added.

Mixtures were incubated in a BioRad T100 thermal cycler: 30 min at 30 °C, then 20 min at 42 °C. Transformations were briefly centrifuged (accelerated to 4,000g, then immediately decelerated without braking). Pellets were washed by resuspension in 50 µL water and re-pelleting. Cells were resuspended in 200 µL C−Trp−Ura medium with 2% glucose and transferred to 96-well culture plates. Cultures were incubated for ∼40 h at 30 °C with shaking. Cultures were either stored (1:1 with 50% glycerol at −80 °C) or induced by 8-fold dilution into SGCAA medium with 0.2% glucose and incubation at 30 °C for 16–24 h with shaking.

#### Yeast display binding assays

20 µL of SGCAA cultures were prepared as described above, and distributed into Corning® 96-well V-bottom plates. Cells were washed with PBSF (PBS + 1% BSA), and then resuspended in 30 µL of 1 µM biotinylated dsDNA and incubated for 30 min at room temperature with shaking. After PBSF washing, cells were stained in 30 µL solution (32 µg/mL anti-c-Myc-FITC, ICL Lab; 32 µg/mL streptavidin-PE, Thermo Fisher, in PBSF) for 20 min at room temperature with shaking. Cells were then washed and resuspended in 200 µL PBSF. Binding was assessed by flow cytometry on an Attune NxT with an autosampler. Data were analyzed with custom Python code and CytoFlow, as in Glasscock et al.^38^. Expression gating used the CytoFlow Gaussian mixture model, and binding was quantified as the PE/FITC intensity ratio for gated events. Binding was quantified as the PE/FITC ratio, using streptavidin-PE to detect biotinylated dsDNA and anti-c-Myc-FITC to detect C-terminal tags.

#### Bacterial expression and purification of DBP fusions

DBP fusion sequences were codon-optimized for *E. coli*, synthesized as G-blocks, and cloned into plasmid LM627 (Addgene #191551) by Golden Gate. Plasmids were transformed into BL21(DE3) *E. coli*. Transformants were inoculated into 2–10 mL LB + 50 mg/L kanamycin starter cultures and incubated overnight at 37 °C, 225 rpm. Starter cultures were diluted 1:50 into 50 mL or 1 L LB + kanamycin and grown at 37 °C, 225 rpm to OD600 0.6–0.8. Expression was induced with 1 mM IPTG, and cultures were grown for ∼16 h at 18 °C, 225 rpm. Cells were harvested by centrifugation at 3,000g for 15 min. Pellets were resuspended in high-salt lysis buffer (2 M NaCl, 20 mM Tris-HCl, EDTA-free protease inhibitor) and sonicated (QSonica Q500, 4-pronged horn, 5 min, 70%). Lysates were clarified (14,000g, 30 min) and supernatants were passed over Ni-NTA resin in gravity columns. Resin was washed with 20 CV of high-salt buffer (2 M NaCl, 20 mM Tris-HCl, 30 mM imidazole, pH 8.0). Proteins were eluted with 2 CV high-salt buffer (2 M NaCl, 20 mM Tris-HCl, 300 mM imidazole) or, for crystallography, cleaved on-column to remove the SNAC-His tag. For cleavage, columns were washed with 5 CV SNAC buffer (100 mM CHES, 100 mM acetone oxime, 100 mM NaCl, 500 mM GnCl, pH 8.6), incubated overnight at room temperature in 5 CV buffer + 0.2 mM NiCl₂, and flowthrough collected. Cleaved or uncleaved eluates were concentrated to 1 mL, filtered, and injected on a Superdex S75 Increase 10/300 GL column (ÄKTA Pure) at room temperature in SEC buffer. High-salt SEC buffer (2 M NaCl, 20 mM Tris-HCl, pH 8.0) was used for crystallography samples and mid-salt buffer (500 mM NaCl, 20 mM Tris-HCl, pH 8.0) for others. Monodisperse fractions were pooled, concentrated with 3 kDa cutoff spin filters (Amicon, Millipore Sigma), and stored at 4 °C. Protein concentrations were determined by A280 on a NanoDrop spectrometer (Thermo Fisher Scientific).

#### Assembly of semi-symmetric protein-DNA complexes

Single-stranded DNA (ssDNA; 97 nt) was purchased from Integrated DNA Technologies (IDT). Equimolar strands were mixed, heated to 72 °C, and slow-cooled to 12 °C over 23 h to form strand-exchanging heterodimers or heterotrimers. Annealed complexes were size-purified by HPLC, collecting fractions at expected UV peaks. Purified DNA complexes were mixed with their associated DBP-fusion proteins to form final protein-DNA complexes, which were again resolved by HPLC to isolate full assemblies, which were then characterized using nsEM.

#### nsEM characterization

For each sample (RNA pseudocycles and protein-DNA complexes), glow-discharged Lacey Carbon copper grids (1 µm holes, 5 µm spacing) received 4 µL diluted RNA, incubated 20 s, blotted, then rinsed with 4 µL nuclease-free H₂O and blotted. Grids were stained three times with 4 µL of 2% uranyl formate, each for 10 s before blotting. Grids dried for ≥5 min before loading into a Talos L120C microscope. Imaging used 120 kV, 2.7 mm Cs, and a Ceta 4K CCD (4096×4096). Automated collection was done with EPU (Thermo Fisher). Data was processed in cryoSPARC to create 2D class averages and 3D reconstructed volumes^48^.

#### Cryo-EM sample preparation

3 µL of purified RNA was applied to glow-discharged 400-mesh ultrathin lacey carbon grids (EMS LC400-Cu-CC-25). Grids were vitrified by plunge-freezing in liquid ethane. Grids were clipped and kept in liquid nitrogen until loading.

#### Cryo-EM data processing

For RNA pseudocycle pC2-240-04, data were collected on a 300 kV FEI Titan Krios. Movies were acquired in SerialEM using beam shifts, at 0.83 Å/pixel. Data were processed in CryoSPARC v4.7^48^. Iterative rounds of particle picking, extraction (400 px box size), and 2D classification were performed. Iterative rounds of ab initio reconstruction were performed (3-15 classes per round), where particle sets associated with volumes of the correct size and shape were selected and fed into subsequent rounds of ab initio reconstruction. Once helical pitch and hairpin extensions became apparent, heterogeneous refinement was performed to remove junk particles, while the target class and its particles were then fed into non-uniform refinement.

#### Cryo-EM model building

The design model was rigid-body docked into the final cryo-EM map in ChimeraX, and backbone-refined to fit the experimental density using ISOLDE^49^. Final RNA structure models were built by iterative map-fitting MD and partial diffusion, where initial fits from ISOLDE were refined by running RFDpoly partial diffusion/denoising to yield cleaner backbone geometry while introducing subtle structural diversification. Three iterations of this process were performed, followed by quick PyRosetta refinement, to produce the final experimentally refined model.

#### Comparison of Cryo-EM model against native RNA folds

To assess whether the designed RNA pseudocycle pC2-240-04 adopts a fold similar to known RNAs, we compared its experimentally-built cryo-EM model against all *representative* RNA structures in the RNAsolo database^50^ (4,089 structures). Structural alignments were performed using USalign^33^ in RNA mode, with pC2-240-04 as the aligned structure and native RNAs as the reference targets. Candidate matches were ranked by TM-score normalized to the design length, and only alignments with ≥0.5 coverage (coverage = *L*_aligned_ / *L*_design_) were considered. The closest structural match was chain-A of the plant mitochondrial ribosome (PDB 6XYW; *L*_aligned_ = 133, *L*_design_ = 240, *L*_native_ = 2,842), which achieved a TM-score of 0.329, supporting the novelty of our *de novo* designed RNA fold.

### IV. Data Availability & Code

The code and weights for RFDpoly will be made available on GitHub at the time of publication. Additionally, tutorial notebooks for carrying out every design campaign described in this manuscript (and more) will be posted online at the time of publication, to demonstrate the use of all features described herein.

Data for OpenKnot designs are available at: https://github.com/eternagame/OpenKnotAIDesignData

## References

[1] Sefah, Kwame, Dihua Shangguan, Xiangling Xiong, Meghan B. O’Donoghue, and Weihong Tan. 2010. “Development of DNA Aptamers Using Cell-SELEX.” Nature Protocols 5 (6): 1169–85.

[2] Kyung-Nam Kang and Yoon-Sik Lee. RNA aptamers: a review of recent trends and applications. Future Trends in Biotechnology, pages 153–169, 2012.

[3] Lee, Jeehyung, Wipapat Kladwang, Minjae Lee, Daniel Cantu, Martin Azizyan, Hanjoo Kim, Alex Limpaecher, et al. 2014. “RNA Design Rules from a Massive Open Laboratory.” Proceedings of the National Academy of Sciences 111 (6): 2122–27.

[4] Zadeh, Joseph N., Conrad D. Steenberg, Justin S. Bois, Brian R. Wolfe, Marshall B. Pierce, Asif R. Khan, Robert M. Dirks, and Niles A. Pierce. “NUPACK: Analysis and design of nucleic acid systems.” Journal of computational chemistry 32, no. 1 (2011): 170–173.

[5] Poppleton, Erik, Niklas Urbanek, Taniya Chakraborty, Alessandra Griffo, Luca Monari, and Kerstin Göpfrich. 2023. “RNA Origami: Design, Simulation and Application.” *RNA Biology*, July, 510–24.

[6] Zadegan, Reza M., and Michael L. Norton. 2012. “Structural DNA Nanotechnology: From Design to Applications.” International Journal of Molecular Sciences 13 (6): 7149–62.

[7] Rothemund, Paul W. K. 2006. “Folding DNA to Create Nanoscale Shapes and Patterns.” Nature 440 (7082): 297–302.

[8] Seeman, N. C. 1982. “Nucleic Acid Junctions and Lattices.” Journal of Theoretical Biology 99 (2): 237–47.

[9] Geary, Cody, Guido Grossi, Ewan K. S. McRae, Paul W. K. Rothemund, and Ebbe S. Andersen. 2021. “RNA Origami Design Tools Enable Cotranscriptional Folding of Kilobase-Sized Nanoscaffolds.” Nature Chemistry 13 (6): 549–58.

[10] Lu, Xiang-jun, and Wilma K. Olson. 2003. “3DNA: A Software Package for the Analysis, Rebuilding and Visualization of Three-dimensional Nucleic Acid Structures.” Nucleic Acids Research 31 (17): 5108–21.

[11] Li, Shuxiang, Wilma K. Olson, and Xiang-Jun Lu. 2019. “Web 3DNA 2.0 for the Analysis, Visualization, and Modeling of 3D Nucleic Acid Structures.” Nucleic Acids Research 47 (W1): W26–34.

[12] Baek, Minkyung, Frank DiMaio, Ivan Anishchenko, Justas Dauparas, Sergey Ovchinnikov, Gyu Rie Lee, Jue Wang et al. “Accurate prediction of protein structures and interactions using a three-track neural network.” Science 373, no. 6557 (2021): 871–876.

[13] Wang, Jue, Sidney Lisanza, David Juergens, Doug Tischer, Joseph L. Watson, Karla M. Castro, Robert Ragotte et al. “Scaffolding protein functional sites using deep learning.” Science 377, no. 6604 (2022): 387–394.

[14] Watson, Joseph L., David Juergens, Nathaniel R. Bennett, Brian L. Trippe, Jason Yim, Helen E. Eisenach, Woody Ahern, et al. 2023. “De Novo Design of Protein Structure and Function with RFdiffusion.” Nature 620 (7976): 1089–1100.

[15] Krishna, Rohith, Jue Wang, Woody Ahern, Pascal Sturmfels, Preetham Venkatesh, Indrek Kalvet, Gyu Rie Lee et al. “Generalized biomolecular modeling and design with RoseTTAFold All-Atom.” Science 384, no. 6693 (2024): eadl2528.

[16] Baek, Minkyung, Ryan McHugh, Ivan Anishchenko, Hanlun Jiang, David Baker, and Frank DiMaio. “Accurate prediction of protein–nucleic acid complexes using RoseTTAFoldNA.” Nature methods 21, no. 1 (2024): 117–121.

[17] Ma, Runze, Zhongyue Zhang, Zichen Wang, Chenqing Hua, Zhuomin Zhou, Fenglei Cao, Jiahua Rao, and Shuangjia Zheng. “RiboFlow: Conditional De Novo RNA Sequence-Structure Co-Design via Synergistic Flow Matching.“ arXiv preprint arXiv:2503.17007 (2025).

[18] Nori, Divya, and Wengong Jin. “Rnaflow: Rna structure & sequence design via inverse folding-based flow matching.” arXiv preprint arXiv:2405.18768 (2024).

[19] Morehead, Alex, Jeffrey Ruffolo, Aadyot Bhatnagar, and Ali Madani. “Towards joint sequence-structure generation of nucleic acid and protein complexes with SE (3)-discrete diffusion.“ arXiv preprint arXiv:2401.06151 (2023).

[20] Anand, Rishabh, Chaitanya K. Joshi, Alex Morehead, Arian R. Jamasb, Charles Harris, Simon V. Mathis, Kieran Didi, Rex Ying, Bryan Hooi, and Pietro Liò. “Rna-frameflow: Flow matching for de novo 3d rna backbone design.” ArXiv (2025): arXiv-2406.

[21] Tarafder, Sumit, and Debswapna Bhattacharya. “RNAbpFlow: Base pair-augmented SE (3)-flow matching for conditional RNA 3D structure generation.“ bioRxiv (2025).

[22] Kubaney, Andrew, Andrew Favor, Lilian McHugh, Raktim Mitra, Robert Pecoraro, Justas Dauparas, Cameron Glasscock, David Baker. “RNA sequence design and Protein–DNA specificity prediction with NA-MPNN.“ bioRxiv (2025).

[23] Dauparas, Justas, Ivan Anishchenko, Nathaniel Bennett, Hua Bai, Robert J. Ragotte, Lukas F. Milles, Basile IM Wicky, et al. “Robust deep learning–based protein sequence design using ProteinMPNN.“ Science 378, no. 6615 (2022): 49–56.

[24] Dauparas, Justas, Gyu Rie Lee, Robert Pecoraro, Linna An, Ivan Anishchenko, Cameron Glasscock, and David Baker. “Atomic context-conditioned protein sequence design using LigandMPNN.” Nature Methods (2025): 1–7.

[25] Cruz, José Almeida, Marc-Frédérick Blanchet, Michal Boniecki, Janusz M. Bujnicki, Shi-Jie Chen, Song Cao, Rhiju Das et al. “RNA-Puzzles: a CASP-like evaluation of RNA three-dimensional structure prediction.” Rna 18, no. 4 (2012): 610–625.

[26] Eterna. OpenKnot. https://eternagame.org/challenges/11843006, 2025. Eterna challenge.

[27] He, Shujun, Rui Huang, Jill Townley, Rachael C. Kretsch, Thomas G. Karagianes, David BT Cox, Hamish Blair, et al. “Ribonanza: deep learning of RNA structure through dual crowdsourcing.“ bioRxiv (2024).

[28] Kretsch, Rachael C., Alissa M. Hummer, Shujun He, Rongqing Yuan, Jing Zhang, Thomas Karagianes, Qian Cong, Andriy Kryshtafovych, and Rhiju Das. “Assessment of nucleic acid structure prediction in CASP16.” bioRxiv (2025): 2025–05.

29. [29] Boitreaud, Jacques, Jack Dent, Matthew McPartlon, Joshua Meier, Vinicius Reis, Alex Rogozhnikov, and Kevin Wu. “Chai-1: Decoding the molecular interactions of life.“ BioRxiv (2024).

[30] Abramson, Josh, Jonas Adler, Jack Dunger, Richard Evans, Tim Green, Alexander Pritzel, Olaf Ronneberger et al. “Accurate structure prediction of biomolecular interactions with AlphaFold 3.” Nature 630, no. 8016 (2024): 493–500.

[31] Das, Rhiju, Rachael C. Kretsch, Adam J. Simpkin, Thomas Mulvaney, Phillip Pham, Ramya Rangan, Fan Bu et al. “Assessment of three-dimensional RNA structure prediction in CASP15.” *Proteins: Structure*, Function, and Bioinformatics 91, no. 12 (2023): 1747–1770.

[32] Zemla, Adam. “LGA: a method for finding 3D similarities in protein structures.” Nucleic acids research 31, no. 13 (2003): 3370–3374.

[33] Zhang, Chengxin, Morgan Shine, Anna Marie Pyle, and Yang Zhang. “US-align: universal structure alignments of proteins, nucleic acids, and macromolecular complexes.” Nature methods 19, no. 9 (2022): 1109–1115.

[34] Marinus, Tycho, Adam B. Fessler, Craig A. Ogle, and Danny Incarnato. “A novel SHAPE reagent enables the analysis of RNA structure in living cells with unprecedented accuracy.” Nucleic acids research 49, no. 6 (2021): e34–e34.

[35] Wayment-Steele, Hannah K., Wipapat Kladwang, Alexandra I. Strom, Jeehyung Lee, Adrien Treuille, Alex Becka, Eterna Participants, and Rhiju Das. “RNA secondary structure packages evaluated and improved by high-throughput experiments.” Nature methods 19, no. 10 (2022): 1234–1242.

[36] Mortimer, Stefanie A., Cole Trapnell, Sharon Aviran, Lior Pachter, and Julius B. Lucks. “SHAPE–Seq: high-throughput RNA structure analysis.” Current protocols in chemical biology 4, no. 4 (2012): 275–297.

[37] Rhiju Das. Openknotscorematlab. https://github.com/eternagame/OpenKnotScoreMATLAB, 2025. GitHub repository. 1.

[38] Glasscock, Cameron J., Robert J. Pecoraro, Ryan McHugh, Lindsey A. Doyle, Wei Chen, Olivier Boivin, Beau Lonnquist et al. “Computational design of sequence-specific DNA-binding proteins.” Nature Structural & Molecular Biology (2025): 1–10.

[39] Holliday, Robin. “A mechanism for gene conversion in fungi.” Genetics Research 5, no. 2 (1964): 282–304.

[40] Praetorius, Florian, and Hendrik Dietz. “Self-assembly of genetically encoded DNA-protein hybrid nanoscale shapes.” Science 355, no. 6331 (2017): eaam5488.

[41] Joung, J. Keith, and Jeffry D. Sander. “TALENs: a widely applicable technology for targeted genome editing.” Nature reviews Molecular cell biology 14, no. 1 (2013): 49–55.

[42] Bhakta, Mital S., and David J. Segal. “The generation of zinc finger proteins by modular assembly.“ In Engineered Zinc Finger Proteins: Methods and Protocols, pp. 3–30. Totowa, NJ: Humana Press, 2010.

[43] Kwon, Diana. “RNA function follows form-why is it so hard to predict?.” Nature 639, no. 8056 (2025): 1106–1108.

[44] Lauko, Anna, Samuel J. Pellock, Kiera H. Sumida, Ivan Anishchenko, David Juergens, Woody Ahern, Jihun Jeung et al. “Computational design of serine hydrolases.” Science 388, no. 6744 (2025): eadu2454.

[45] Yang, Jianyi, Ivan Anishchenko, Hahnbeom Park, Zhenling Peng, Sergey Ovchinnikov, and David Baker. “Improved protein structure prediction using predicted interresidue orientations.” Proceedings of the National Academy of Sciences 117, no. 3 (2020): 1496–1503.

[46] Tran, Long, Shajesh Sharma, Steffen Klein, David Jurgens, Justin Decarreau, Bingxu Liu, Yujia Wang et al. “Design of Orthogonal Far-Red, Orange and Green Fluorophore-binding Proteins for Multiplex Imaging.” bioRxiv (2025): 2025–08.

[47] Kretsch, Rachael C., Yuan Wu, Svetlana A. Shabalina, Hyunbin Lee, Grace Nye, Eugene V. Koonin, Alex Gao, Wah Chiu, and Rhiju Das. “Naturally ornate RNA-only complexes revealed by cryo-EM.” Nature (2025): 1–3.

[48] Punjani, Ali, John L. Rubinstein, David J. Fleet, and Marcus A. Brubaker. “cryoSPARC: algorithms for rapid unsupervised cryo-EM structure determination.” Nature methods 14, no. 3 (2017): 290–296.

[49] Croll, Tristan Ian. “ISOLDE: a physically realistic environment for model building into low-resolution electron-density maps.” Biological Crystallography 74, no. 6 (2018): 519–530.

[50] Adamczyk, Bartosz, Maciej Antczak, and Marta Szachniuk. “RNAsolo: a repository of cleaned PDB-derived RNA 3D structures.” Bioinformatics 38, no. 14 (2022): 3668–3670.

